# Misalignment between perceived and actual ability on a balance beam walking task

**DOI:** 10.1101/2025.11.10.687691

**Authors:** Sam Beech, Christina Geisler, Sierra Creer, Cecilia Monoli, Anna C. Render, Peter C. Fino

**Affiliations:** University of Utah, Department of Health & Kinesiology, Salt Lake City, UT; University of Utah, Department of Psychology, Salt Lake City, UT

**Keywords:** Balance, Recalibration, Perceived balance

## Abstract

Effective dynamic balance control is necessary to maintain stability, but it is an individual’s self-perceived ability that ultimately determines movement selection. Accurate self-estimation of balance ability is therefore essential to ensure that movement choices align with true capability. This study examined individuals’ perception of their ability on a clinically standardized Narrowing Beam Walking Task (NBWT) to examine 1) the initial perception of balance ability before attempting the task, and 2) how experience completing the task improves the accuracy of self-perceived balance. Collegiate athletes provided self-estimates of performance at baseline (before any trials with the task), early-training (after completing two trials), and post-training (after a further 8 trials). Actual task performance was quantified using the final 8 trials. At baseline, athletes poorly estimated their ability: individuals with poorer task performance tended to overestimate their ability while higher-performing individuals tended to underestimate their ability. With practice, absolute estimation error significantly decreased, indicating that task-specific exposure facilitated recalibration to bring self-estimates of performance in closer alignment to actual performance. These effects were consistent across all tested sporting disciplines. These findings show that effective balance control and frequent engagement in similar, but unrelated balance tasks, does not facilitate accurate self-perception of performance on the NBWT. Instead, brief task-specific exposure was required to refine balance estimates. These findings have implications for balance testing and rehabilitation that seeks to improve mobility in populations whose misjudgments of balance ability are often associated with negative outcomes, such as falls.

## INTRODUCTION

Balance control is a fundamental component of locomotion, allowing individuals to move safely and efficiently through their environments. Lab-based assessments consistently link effective dynamic stability to functional movement (Harrison et al., 2021), reduced injury risk (Emery et al., 2005), and improved quality of life (Dunsky et al., 2019). However, movement in daily life is not determined solely by physical capacity. Rather, action selection is often guided more strongly by perceived ability, with self-estimated capability frequently serving as the better predictor of behavior (Bandura, 1977; Bruce et al., 2012; Morasso et al., 2015). During action, we continuously generate estimates of our capabilities with varying degrees of accuracy (Creem-Regehr et al., 2019; Franchak & Adolph, 2014; Warren, 1984). Therefore, beyond developing effective dynamic balance, maintaining an accurate estimation of self-perceived ability is essential to ensure that movement selection aligns with true capability.

There is growing recognition for the importance of accurate self-perception as balance misjudgments carry meaningful consequences. Approximately one-third of older adults misjudge their balance ability, with errors occurring in both directions (Delbaere et al., 2010). Underestimation promotes risk-averse behaviors and activity restriction, which reduce physical activity and, counterproductively, increase fall risk (Delbaere et al., 2010; Hadjistavropoulos et al., 2011). Overestimation, on the other hand, leads individuals to attempt movements beyond their stability limits, thereby increasing the risk of falls (Kawasaki & Tozawa, 2020; Sakurai et al., 2013). Similar misjudgments of locomotor control and balance that are linked to fall risk have also been observed in individuals with Parkinson’s disease (Kamata et al., 2007; Lafargue et al., 2013; Longhurst et al., 2025), stroke (Morone et al., 2014; Sakai et al., 2024), and moderate-to-severe traumatic brain injury (Hays et al., 2019). Collectively, these findings highlight a need to determine how accurate self-estimates are developed.

Emerging evidence suggests that increased practice within a given task leads to recalibration of inaccurate self-perceptions. Quammen et al. (2025) demonstrated that both expert and non-expert soccer players initially overestimated their ability to intercept a ball, but both groups rapidly recalibrated their judgments with increased task exposure, leading to accurate self-estimates of performance. Ellmers et al. (2018) reported similar recalibration effects for static balance in older adults, where estimated and actual ability became more closely aligned following a four-week Wii balance board intervention. These findings indicate that baseline self-perceptions are often unreliable, even among skilled performers, but that task exposure promotes recalibration.

However, it remains unknown whether these recalibration effects also occur within dynamic balance tasks, which more closely reflect the demands of everyday locomotion. Although dynamic balance training reliably improves objective stability (Lesinski et al., 2015), its impact on the accuracy of self-estimated performance has yet to be established.

The purpose of this study was to determine if people can accurately estimate their ability on a balance beam task – the clinically standardized Narrowing Beam Walking Task (NBWT) (Sawers & Hafner, 2018) – and whether exposure to the task improves the accuracy of self-estimated dynamic balance. Collegiate athletes were recruited as a model of a high-functioning population, as they generally exhibit superior balance relative to non-athletes (Bonis & Tillery, 2021; Davlin, 2004) and have previously shown rapid recalibration in motor tasks (Quammen et al., 2025). Self-estimates of task performance were recorded at three timepoints – baseline, early-training, and post-training – and compared against actual task performance. We hypothesized that individuals would initially overestimate their balance ability at baseline, but that estimation error would decline with repeated task exposure, leading to greater alignment between self-estimated and actual performance in the early-training and post-training timepoints.

We also investigated whether self-estimation accuracy differed across sporting disciplines to determine if expertise in similar tasks (e.g., gymnastics) would transfer to the NBWT. This comparison provided an opportunity to explore whether frequent engagement in related, though not identical, balance-focused activities influences self-estimation accuracy and recalibration.

## MATERIALS and METHODS

### Participants

Fifty-seven healthy young adult athletes participated in this study. This sample included 24 female athletes (Mean age = 19.04, SD = 1.37) and 33 male athletes (Mean age = 19.09, SD = 1.83). Demographic information by athletic discipline is provided in Table 1. Eligibility criteria required participants to be at least 18 years of age, have no history of concussion within the past year, and no recent or planned lower-limb surgery that could interfere with walking or balance. All participants provided written informed consent prior to participation, and all procedures were approved by the Institutional Review Board at the University of Utah.

**Table 1.**
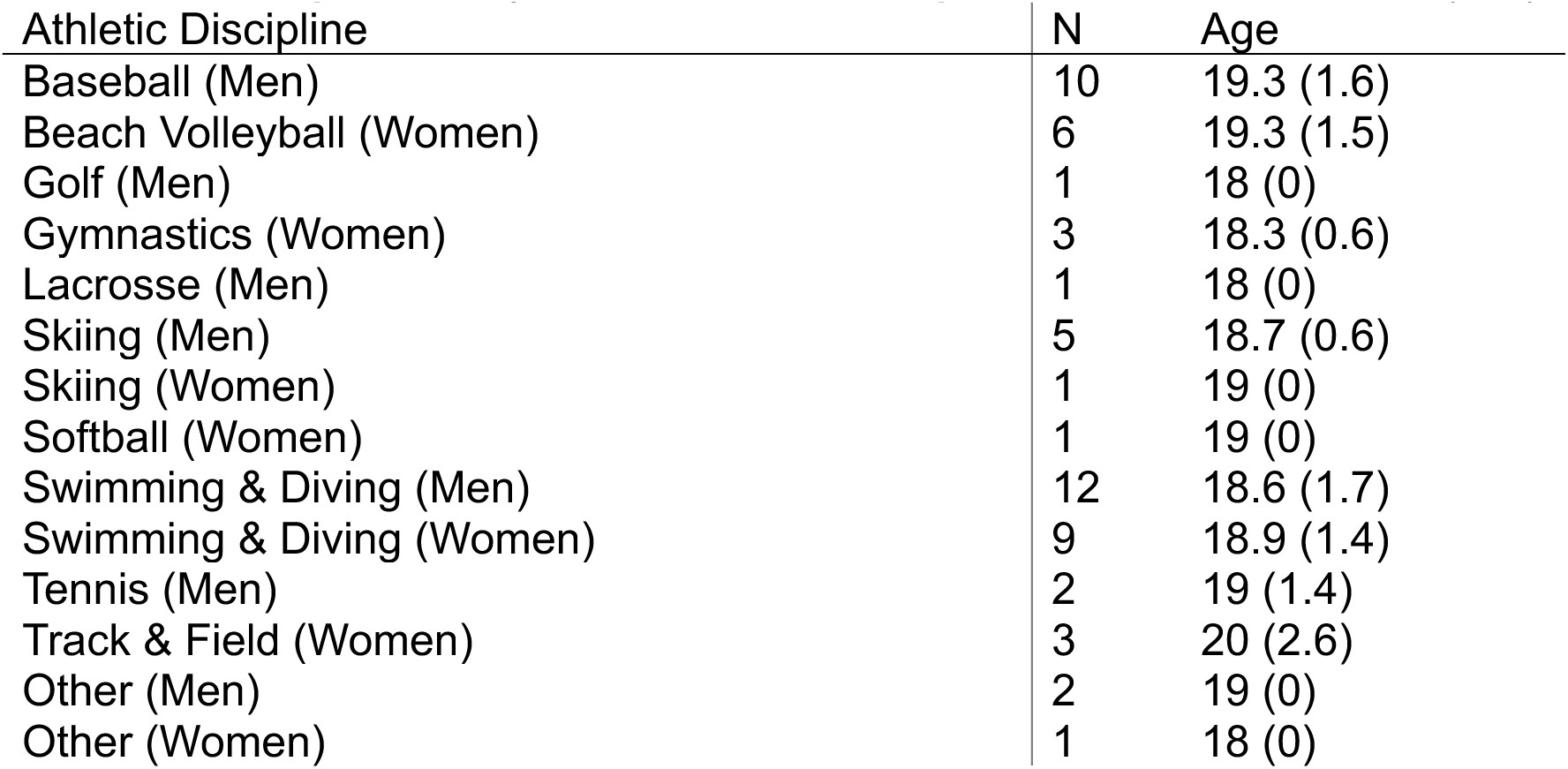
Demographics by athletic discipline. Ages represented as mean (s.d.)

### Procedures and design

A mixed-measures design was employed in which athletes from different sports provided self-estimates of performance at three timepoints while completing the NBWT (Sawers & Hafner, 2018). The task required participants to walk across four progressively narrower beams, labelled A, B, C, and D. Each beam was 1.83 m (6.00 ft) long and 3.81 cm (1.50 in) tall, and beams A, B, C, and D were 18.42 cm (7.25 in), 8.89 cm (3.50 in), 3.81 cm (1.50 in), and 1.91 cm (0.75 in) wide, respectively. Participants wore sneakers and crossed their arms over their chest while attempting the task. To account for the higher level of function in elite athletes compared to the clinical populations for which the NBWT was originally designed (Sawers & Hafner, 2018), participants were instructed to walk heel-to-toe as far along the sequence of connected beams as possible without stepping off. Walking speed was not constrained and was not recorded. A trial ended when the participant’s foot touched the floor, they uncrossed their arms, or they successfully walked the full length of the beams. The completed distance for each trial was recorded based on the heel position of the trailing limb, with a maximum distance of 6.71m (22 ft) – excluding the first 0.61 m (2 ft) of beam A to allow for one full step (Sawers & Hafner, 2018). A beam was considered successfully completed when both heels crossed its end. The participants completed ten trials in total, but the first two were discarded as practice trials. Actual ability was recorded as the number of times out of 8 they completed each beam in trials 3 – 10.

Self-perceived ability was assessed at three points: baseline – before completing any trials; early-training – after completing the two practice trials; and post-training – after completing the eight experimental walking trials. At each timepoint, participants were asked, “Before you begin, we would like you to estimate how far you think you can walk on this beam. Out of 8 trials, how many times do you think you could make it completely past beam [A/B/C/D]”, for each of the four beams.

### Data Analysis

The completion of each beam was converted into corresponding distance values to better reflect progression through the task: non-completion of beam A was coded as 0m (0 ft); completion of beam A as 1.22m (4 ft); and completion of beams B, C, and D as 3.05m (10 ft), 4.88m (16 ft), and 6.71m (22 ft), respectively. Success (1) and failure (0) were coded as binary outcomes, providing a maximum score of 8 for each beam based on 8 scored trials. These binary data were used to fit logistic regression models for each participant and timepoint using MATLAB’s ‘glmfit’ function with a binomial distribution and logit link. In accordance with Quammen et al. (2025), we extracted the chance threshold from each model, defined as the distance corresponding to a 50% predicted success rate. In cases where participants estimated, or completed all four beams, this chance threshold was set to the maximum distance – 6.71m (22 ft).

We calculated the direction and magnitude of the estimation error by comparing the signed difference between each participant’s self-estimated 50% threshold and their actual 50% performance threshold at the three estimation timepoints (baseline, early-training, post-training). The value was the error in meters, while the sign represented the direction of error: positive values reflected overestimation and negative values reflected underestimation. The absolute estimate error was recorded to identify the magnitude of error regardless of direction. We compared the directional estimate error and absolute estimate error at the three self-estimation timepoints using Linear Mixed Effects Modelling (LMM) with a Satterthwaite approximation and F-test for fixed effects. The model included the timepoint, the participant’s sport discipline, and gender as categorical fixed effects. The participant number was also included as a random effects factor with random intercepts and slopes for the repeated ‘timepoint’ factor.

We also examined the relationship between actual balance ability and baseline estimation accuracy using a Pearson’s correlation between each participant’s 50% performance threshold and their directional estimation error at baseline. All analyses were completed in JASP using a significance level of 0.05 for all comparisons (Version 0.19.1; University of Amsterdam, 2024).

## RESULTS

On average, people underestimated their balance ability at baseline and early training, followed by overestimation at post-training (Table 2). However, this directional estimation error varied widely across timepoint and by individual, with some participants largely underestimating and others overestimating their balance ability at baseline, and changing between early training, and post-training (Figure 1). The main effect of the estimation timepoint was significant, (F(2, 112) = 9.862, p < .001) (Table 2). Post-training directional estimate error was significantly larger than at baseline, (β = 0.506, t(112) = 3.863, p < .001), and early-training (β = 0.502, t(112) = 3.829, p < .001), while there was no significant difference between baseline and early-training, (β = 0.004, t(112) = 0.033, p = .974). There was no effect of sport discipline (F(12, 43) = 0.864, p = .587) and no effect of gender (F(1, 43) = 2.632, p = .112).

**Figure 1.**
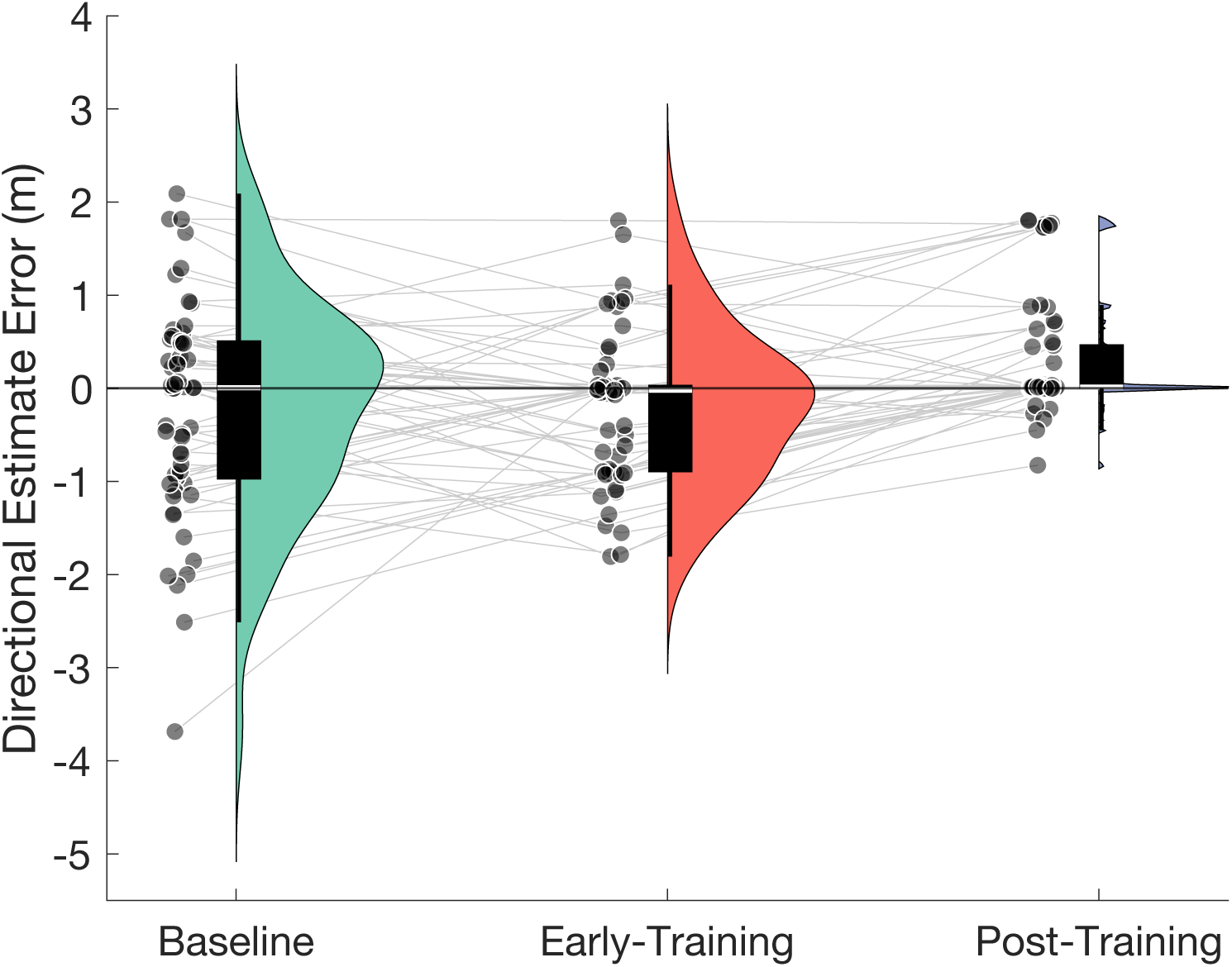
Directional estimate error across each timepoint. Negative values reflect underestimation and positive values reflect overestimation. Scattered points are individual subjects, with thin gray lines connecting timepoints within a subject. Violin plots illustrate the distribution, with box plots showing the median (white line) and quartile ranges (black box).

**Table 2.** The estimated marginal means and confidence intervals (CI) for directional and absolute estimate error.

| Error Type | Timepoint | Estimated Marginal Mean | Lower 95% CI | Upper 95% CI |
| --- | --- | --- | --- | --- |
| **Directional** | Baseline | -0.378 | -0.673 | -0.084 |
|  | Early-training | -0.374 | -0.668 | -0.080 |
|  | Post-training | 0.128 | -0.166 | 0.422 |
| **Absolute** | Baseline | 0.856 | 0.653 | 1.058 |
|  | Early-training | 0.578 | 0.376 | 0.781 |
|  | Post-training | 0.348 | 0.146 | 0.551 |

Completion of the NBWT significantly improved the alignment between estimated and actual performance. Absolute estimate error significantly decreased with training (F(2, 112) = 11.077, p < .001; Figure 2; Table 2). Post-training absolute error was smaller than both baseline (β = -0.507, t(112) = -4.700 p < .001) and early-training (β = --0.230, t(112) = -2.131, p = .035). Early-training absolute error was also smaller than baseline (β = –0.277, t(112) = –2.569, p = .023). There was no effect of sport discipline (F(12, 43) = 1.235, p = .292) and no effect of gender (F(1, 43) = 0.697, p = .409).

**Figure 2.**
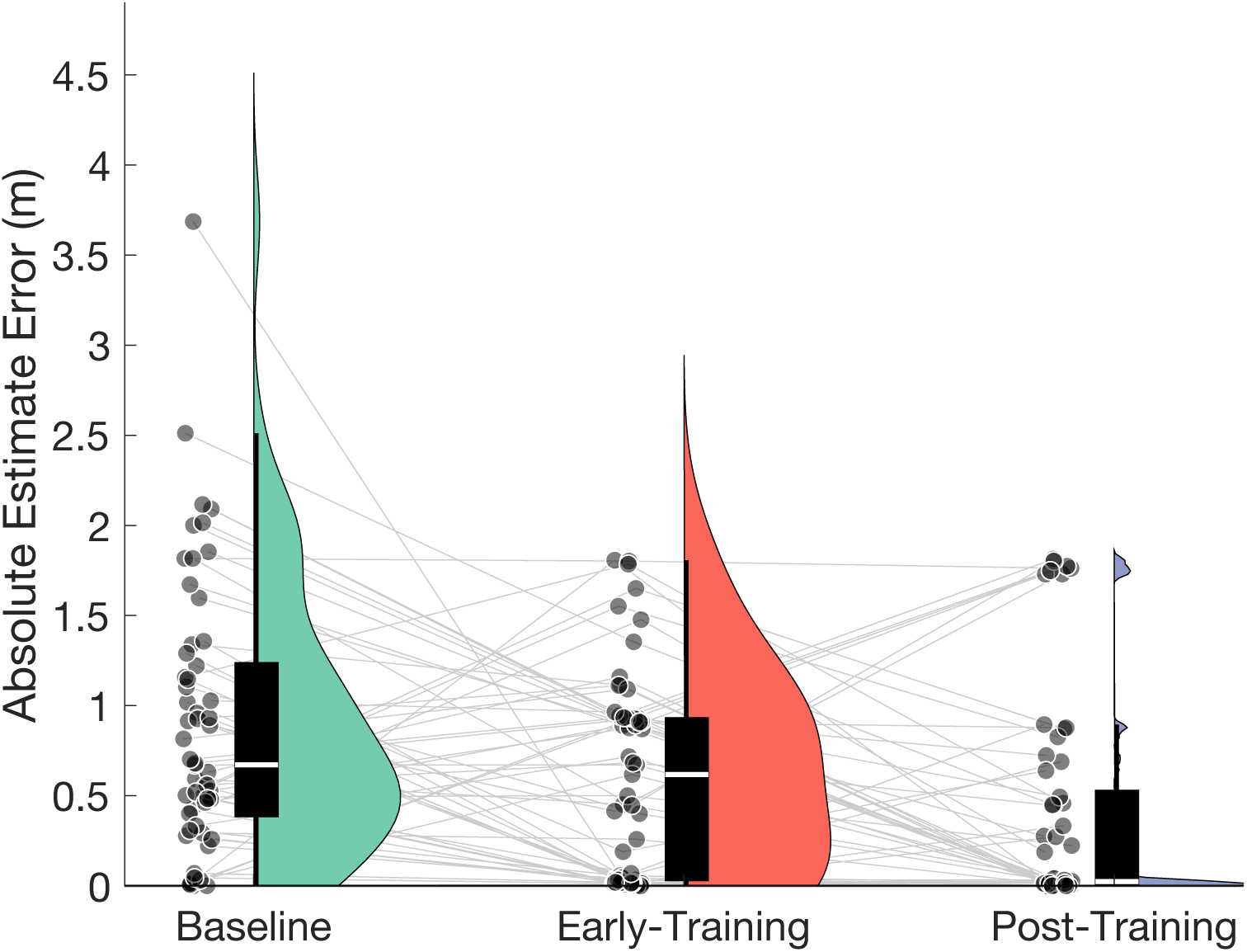
Absolute estimate error across each timepoint. Individuals significantly decrease absolute error from baseline to post-training. Scattered points are individual subjects, with thin gray lines connecting timepoints within a subject. Violin plots illustrate the distribution, with box plots showing the median (white line) and quartile ranges (black box).

Baseline directional error was moderately correlated with actual NBWT performance (r(55) = –.556, p < .001, R² = .309; Figure 3). Participants with poorer balance tended to overestimate their ability and those with better balance tended to underestimate their ability.

**Figure 3.**
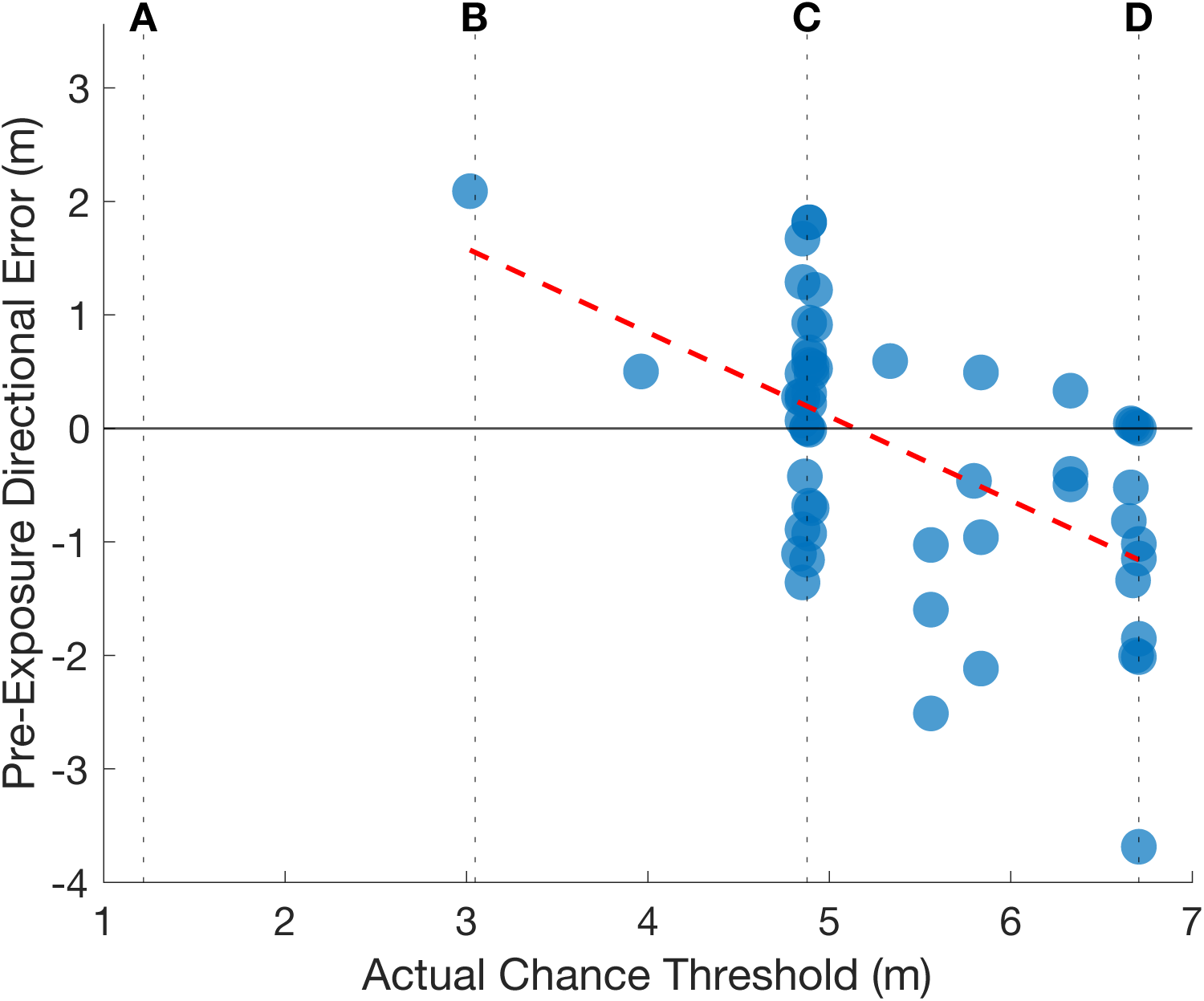
Relationship between baseline directional error estimate and actual performance. Positive and negative values reflect overestimation and underestimation, respectively. The beam numbers are placed over their corresponding distance. People who have better balance ability tend to underestimate their ability, while people with relatively worse balance ability tend to overestimate their ability.

## DISCUSSION

This study investigated whether people can accurately estimate their ability on a balance beam task (the NBWT), and whether brief practice on the NBWT would improve the accuracy of self-estimated dynamic balance ability. Contrary to our hypothesis, participants, on average, tended to underestimate their performance at baseline, but the large variability showed no consistent directional bias. As expected, absolute estimate error decreased following training, showing that practice on the NBWT promoted recalibration and brought self-perceived ability into closer alignment with actual performance. These patterns were consistent between genders and across all sport disciplines.

The improved self-estimation accuracy is consistent with previous recalibration research. Quammen et al. (2025), reported that estimation error for successful interception of a soccer ball decreases with increased task exposure. Furthermore, older adults display improved self-estimation accuracy in static balance control following a four-week Wii balance board intervention (Ellmers et al., 2018). The present results complement this existing literature by showing that a clinically accessible NBWT effectively improves the accuracy of self-perceived dynamic balance ability. Notably, we observed a significant reduction in the absolute estimation error after just two practice trials, indicating rapid recalibration of self-perceived balance ability. This finding demonstrates that even brief exposure can elicit meaningful cognitive recalibration of perceived stability, emphasizing the potential of short, targeted practice for refining self-estimation accuracy.

The observation of recalibration in our collegiate athlete population underscores the importance of task-specific training in correcting self-perceived errors in balance control. This finding is particularly striking given that athletes not only demonstrate superior balance control (Bonis & Tillery, 2021) but are also expected to possess precise internal models of their motor capabilities developed through extensive practice and experience (Du et al., 2022; Yarrow et al., 2009). Despite these advantages, they were initially poor at estimating their dynamic balance ability, but rapidly recalibrated and improved with increased practice on the NBWT. Importantly, baseline estimation accuracy was not aligned with actual performance: athletes with poorer balance tended to overestimate, whereas those with better balance tended to underestimate. Moreover, athletes across all sporting disciplines displayed similar initial errors and recalibration patterns. These results parallel findings from balance training research showing that learning is highly task-specific and does not generalize to similar tasks (Giboin et al., 2015). Extending this principle to self-perception, our findings demonstrate that accurate self-estimation does not emerge from superior balance capacity or generalized exposure to balance-focused activities but instead requires task-specific experience.

The observation of task-specific recalibration carries important clinical and functional implications. Misjudgments of balance associated with fall risk have been observed in older adults (Delbaere et al., 2010; Hadjistavropoulos et al., 2011; Kawasaki & Tozawa, 2020; Sakurai et al., 2013) and in clinical populations with Parkinson’s disease (Kamata et al., 2007; Lafargue et al., 2013; Longhurst et al., 2025), stroke (Morone et al., 2014; Sakai et al., 2024), and moderate-to-severe traumatic brain injury (Hays et al., 2019). While conventional balance training improves objective stability within these populations (Conradsson et al., 2015; Fritz & Basso, 2013; Lesinski et al., 2015; Yavuzer et al., 2006), our results suggest recalibration may complement physiological improvements to promote safer and more confident mobility. More accurate estimates of one’s ability could, for instance, enhance balance confidence and encourage more active behavior independent of physical capacity (Danks et al., 2016; French et al., 2015; Miller et al., 2022). However, the specificity of the movement must be considered. Balance misjudgments in a ‘stepping down’ task did not significantly predict falling in older adults (Kluft et al., 2019), potentially because the ‘stepping down’ movement does not generalize to other movements in which falls commonly occur.

Additionally, initial misjudgments of self-perceived locomotor stability do not correlate between tasks (Kluft et al., 2017). Therefore, like balance training, it is important to identify specific movements in which falls often occur as targets for recalibration.

Importantly, the brief period of exposure required for recalibration observed in our participants indicates that multiple movements could be targeted within a single session.

It remains unclear whether the recalibration observed here generalizes beyond the NBWT to these broader, ecologically relevant contexts. The absence of differences between athletes across sporting disciplines suggests that recalibration may not generalize to other tasks. However, it is important to consider that our participants were young, healthy athletes whose estimation errors carried minimal consequences for daily function. In contrast, for older adults and clinical populations, misjudgments of balance have real-world consequences for both physical safety and psychological wellbeing. In these groups, even task-specific practice in controlled settings may yield broader benefits: individuals who underestimate may gain confidence to engage in safe activities they previously avoided, whereas those who overestimate may learn to move with greater caution. Furthermore, such psychological recalibration could transfer to other situations. The extent to which recalibration of self-perceived balance on a clinically standardized, lab-based balance task generalizes to ecologically relevant tasks such as stair negotiation, obstacle crossing, or complex gait challenges, remains currently unknown.

## CONCLUSION

At initial exposure, athletes across all sporting disciplines demonstrated poor accuracy in estimating their performance on the NBWT. Misjudgments at baseline spanned the entire ability spectrum, with poorer performers tending to overestimate and better performers tending to underestimate their ability. However, after 10 trials, participants across all sporting disciplines showed reduced estimation error. These findings demonstrate that better actual balance ability does not ensure accurate self-estimates. Instead, task-specific exposure is required to recalibrate self-perceptions of balance, bringing estimated and actual ability into closer alignment.

## ACKNOWLEDGEMENTS

The authors would like to thank Zosia Welgarz, Kelsi Schiltz, and Noemi Montemurro for assisting in data collection, as well as the University of Utah Athletic Training staff for assisting in participant recruitment.

